# A Bayesian Approach for Identifying Driver Mutations within Oncogenic Pathways through Mutual Exclusivity

**DOI:** 10.1101/2025.05.27.656485

**Authors:** Xinjun Wang, Caroline Kostrzewa, Allison Reiner, Ronglai Shen, Colin Begg

**Affiliations:** Department of Epidemiology and Biostatistics, Memorial Sloan Kettering Cancer Center, New York, NY, USA

**Author notes:** Corresponding authors: Xinjun Wang and Colin Begg.

## Abstract

Distinguishing driver mutations from the large background of passenger mutations remains a major challenge in cancer genomics. Evidence-based approaches to nominate driver mutations are often limited by the availability of experimental or clinical validation for specific variants. As clinical sequencing becomes integrated into patient care, computational methods provide powerful opportunities to analyze expanding genomic datasets and identify functional candidates beyond the current knowledge base. Among various analytical frameworks, mutual exclusivity, the observation that mutations in two or more genes tend not to co-occur within the same tumor, has been particularly attractive. Building on this principle, we propose BayesMAGPIE, a refined version of a statistical method, MAGPIE, developed previously for identifying driver genes within oncogenic pathways. The new method introduces two key innovations. First, it incorporates information on mutation type using a Bayesian hierarchical modeling framework, enabling the distinction between potential differences in functional effects among variants within the same gene, thereby improving the accuracy of driver identification. Second, it models gene-specific driver frequencies with a Dirichlet prior which effectively controls the sparsity of the inferred driver set and aligns with the biological expectation that most tumor types are driven by a small number of genes. We evaluate BayesMAGPIE through extensive simulation studies to assess its estimation bias and accuracy in driver identification, and benchmark its performance against MAGPIE using TCGA data from eight cancer types.

## 1 INTRODUCTION

Cancer is a genetic disease driven by the accumulation of somatic mutations. Tumor genomic sequencing often reveals numerous mutations, presenting a key challenge of distinguishing driver mutations, those critical to tumor initiation and progression, from passenger mutations. A number of computational methods have been developed to identify driver mutations, broadly categorized by their underlying strategy. Frequency-based methods such as MutSigCV (Lawrence et al., 2013), DriverML (Han et al., 2019), ActiveDriver (Reimand and Bader, 2013), and OncodriveFML (Mularoni et al., 2016) identify genes with mutation rates exceeding background expectations. Hotspot-based approaches such as OncoDriverCLUST (Tamborero, Gonzalez-Perez and Lopez-Bigas, 2013) and MSEA (Jia et al., 2014) detect gain-of-function mutations within specific protein domains. Network-based methods, including DawnRank (Hou and Ma, 2014) and DriverNet (Bashashati et al., 2012), leverage known biological interactions to identify clusters of driver genes.

Another influential concept is mutual exclusivity, the observation that mutations in two (or more) genes tend not to occur together in the same tumor, suggesting functional redundancy or synthetic lethality within oncogenic pathways. Important examples of mutual exclusivity include *EGFR* and *KRAS* in lung adenocarcinoma (Ding et al., 2008; Network, 2014; Pao et al., 2005), and *BRAF, NRAS*, and *NF1* in the RTK-RAS pathway in melanoma (Cancer Genome Atlas, 2015; Shoushtari et al., 2021). Many statistical methods have been developed to detect mutually exclusive gene sets, including greedy search algorithms based on novel criteria or score metrics (Dendrix (Vandin, Upfal and Raphael, 2012) and Mutex (Babur et al., 2015)), graph-based approaches (MEMo (Ciriello, Cerami, Sander and Schultz, 2012) and gcMECM (Hu, Yan, Chen and Meerzaman, 2021)), computational-oriented methods that can scale up to genome-wide analysis (WeSME (Kim, Madan and Przytycka, 2017) and FaME (Fedrizzi et al., 2021)), and permutation-based tests (CoMEt (Leiserson, Wu, Vandin and Raphael, 2015) and WExT (Leiserson, Reyna and Raphael, 2016)). Additionally, MEGSA (Hua et al., 2016), DISCOVER (Canisius, Martens and Wessels, 2016), TiMEx (Constantinescu et al., 2016), MEScan (Liu et al., 2021) and a method proposed by Szczurek et al. (2014) evaluate mutual exclusivity using formal statistical inference.

While most existing methods focus on de novo discovery of mutually exclusive gene sets, our proposed approach leverages mutual exclusivity within predefined oncogenic pathways, i.e., gene sets that are known to share functional roles (Sanchez-Vega et al., 2018). Within each pathway, we aim to identify probabilistically the specific genes within a pathway and the individual variants within such genes that are mostly likely the true drivers. A key modeling assumption is that, within any given pathway, a tumor can harbor at most one driver mutation though additional drivers may exist in other pathways.

We present our proposed method as an advancement over our earlier work, MAGPIE (Wang et al., 2024), which was built upon the likelihood framework of MEGSA, developed by Hua et al. (2016). In both the MAGPIE and MEGSA frameworks, a mutation in a given gene can represent either a driver or a passenger event. Passenger mutations in this context refer to random, non-functional somatic mutations arising from the genetic instability commonly observed in tumor cells. These models leverage global patterns of mutual exclusivity across tumors to estimate the proportion of samples harboring a driver mutation within a given pathway. This is based on the key assumption of mutual exclusivity, i.e., a tumor can harbor at most one driver mutation in the pathway under investigation. Our analysis is limited to non-synonymous mutations, except when calculating tumor mutational burden (TMB), which serves as a confounding factor.

In our new approach, we introduce two major innovations not incorporated in previous methods. First, we integrate mutation-type information (e.g., missense vs. truncating) into the model in recognition that functionality in a specific gene may be largely restricted to one of these sub-types. Second, we adopt a Bayesian hierarchical modeling framework to account for the more complex structure of the data. This modeling framework allows us to incorporate gene-specific driver frequencies using a Dirichlet prior, which helps control the sparsity of inferred driver patterns through tunable hyperparameters. Ultimately, our method enables probabilistic classification of each mutation in each tumor as either a driver or a passenger, guided by the global empirical patterns of mutual exclusivity in the data.

In summary, our proposed method provides a comprehensive probabilistic framework for identifying genes with strong evidence of being drivers and pinpointing the specific mutations within those genes that are most likely driving tumorigenesis. Through a detailed analysis of the RTK-RAS pathway in the Cancer Genome Atlas Lung Adenocarcinoma (TCGA-LUAD) dataset, we demonstrate that our method successfully identifies well-known recurrent mutations and also highlights rare, potentially important mutations. To evaluate performance, we benchmark our method against MAGPIE using TCGA data from seven additional cancer types, offering a broad assessment under real-world conditions. We further conduct extensive simulation studies to assess the accuracy of our method and to provide practical guidance on prior selection. Our method is implemented in a Python package named BayesMAGPIE (<u>Bayes</u>ian <u>m</u>utual exclusivity <u>a</u>nalysis of cancer <u>g</u>enes and variants and their <u>p</u>robability of being driv<u>e</u>r).

## 2 METHOD

Our analytic framework is pathway-specific, and the analysis is restricted to a set of *m* genes in the pathway under consideration. A key assumption of the model is that although a tumor may harbor multiple driver mutations overall, it can have at most one driver mutation within the specific pathway being analyzed.

### 2.1 MAGPIE Framework

Our new model strategy is constructed based on an existing MAGPIE framework developed by Wang et al. (2024). Let *x*_*i*_ = (*x*_*i*1_, … , *x*_*im*_) denote the observed binary mutation status of tumor *i* (*i* = 1, … , *n*), where *x*_*ij*_ = 1 if a non-synonymous alteration is observed in gene *j* (*j* = 1, … , *m*) and 0 otherwise. Any observed mutation must be either a driver mutation or a passenger mutation, and only one true driver mutation from the pathway is possible in a given tumor. As a result, all true driver mutations observed in the tumor must be mutually exclusive, i.e., no two mutations in a given tumor can be functioning as drivers simultaneously. If two occur, one must simply be functioning as an additional, passenger mutation in this particular tumor.

Assume that the gene membership of the driver mutation for tumor *i* in the study cohort is denoted by 𝒛_*i*_ = (𝑧_*i*0_, 𝑧_*i*1_, … , 𝑧_*im*_), a latent variable to be inferred. Tumors with 𝑧_*i*𝑘_ = 1, 𝑘 > 0 have driver mutations in the 𝑘^𝑡ℎ^ gene, while tumors with 𝑧_*i*0_ = 1 do not possess a driver mutation in the pathway. Let 𝝉 = (𝜏_0_, 𝜏_1_, … , 𝜏_*m*_) denote the vector of mixture proportions of tumors having each gene-specific driver mutation, i.e., 𝜏_𝑘_ = 𝑝(𝑧_*i*𝑘_ = 1). Each gene is also assumed to have a constant passenger mutation rate, denoted by 𝜋_*j*_ for the *j*^𝑡ℎ^gene, which is independent of driver mutations. Under a standard mixture model framework, the log likelihood of the observed data is given by

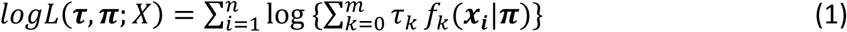

where 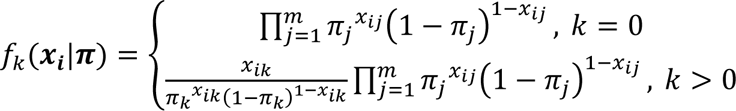

is the cluster-specific probability density function of *x*_*i*_.

Global covariates (i.e., unconditional on the existence of any specific observed mutation) such as tumor mutational burden (TMB) can be adjusted in the model. Let 𝑦_*i*_ denote the TMB score for tumor *i*, defined as the centered logarithm of the total number of observed mutations (including synonymous mutations) across all sequenced genes. To account for the influence of TMB on background passenger mutations within each tumor, the tumor-specific passenger mutation rates are identified using {𝜋_*ij*_} rather than {𝜋_*j*_}. Given 𝜋_*ij*_ is bounded by 0 and 1, the covariate adjustment is modeld by

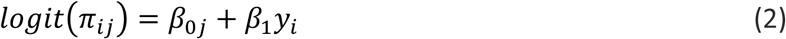

where 𝛽_0*j*_ represents the baseline log odds of the passenger mutation rate for the *j*^𝑡ℎ^ gene, and 𝛽_1_captures the common effect of TMB on the passenger mutation rate across all genes in the pathway. The log likelihood of the observed data adjusting for TMB is then given by

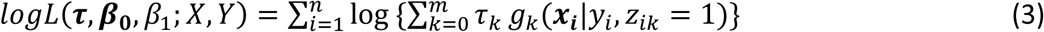

where

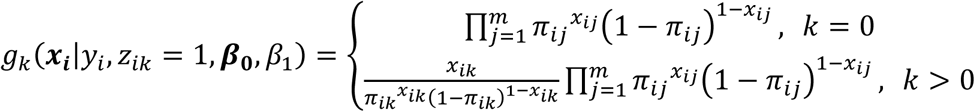

is the cluster-specific probability density function of *x*_*i*_ conditional on 𝑦_*i*_, and 𝜋_*ij*_ = 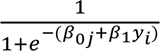.

MAGPIE identifies driver mutations by leveraging patterns of mutual exclusivity across tumors. For each tumor, the goal is to determine which gene harbors the driver mutation within a pre-specified pathway, or to infer that no driver mutation is present in the pathway. This inference is performed probabilistically using posterior probabilities derived via Bayes’ rule. Specifically, the posterior probability that the mutation in the 𝑘^𝑡ℎ^ gene (𝑘 > 0) in the pathway is the driver in the *i*^𝑡ℎ^ tumor is

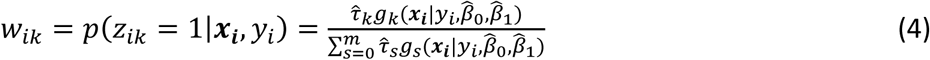

When 𝑘 = 0, equation (4) provides the posterior probability that tumor *i* has no driver mutation in the pathway. In cases where a tumor is observed to have mutations in multiple genes, the most likely driver gene is identified by selecting the index 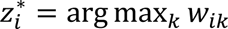.

### 2.2 Proposed Approach – BayesMAGPIE

The leverage used in MAGPIE for distinguishing driver genes is contained in the broad patterns of mutual exclusivity observed. However, mutual exclusivity patterns can only provide direct evidence for entities that occur frequently, such as somatic mutational patterns in entire genes, or in the very few individual hotspot variants. The new approach is motivated by the belief that we can extract more precise evidence about individual driver events by incorporating covariates that contain variant-level information into a statistical model.

We first note that covariates can affect the mutation probabilities in two distinct ways: by affecting the background passenger mutation rate, denoted 𝜋_*ij*_ for the *j*^𝑡ℎ^ gene in the *i*^𝑡ℎ^ tumor, or by directly affecting the probability that an observed mutation is a driver mutation, denoted 𝜏_*j*_. In MAGPIE, factors such as TMB that can influence the background passenger mutation rate are adjusted using a regression model that is presumed to affect all genes in a proportional manner, but without directly influencing the relative frequency of drivers. In the new approach we seek to create a modeling structure to permit the use of factors that could have a strong influence on whether or not an observed mutation is a driver. Examples of such kinds of factors are those genetic factors that are known to affect the functionality of individual variant types. A factor likely to have considerable importance is the mutation type: e.g., missense vs nonsense vs frameshift vs deletion. It is also likely that the effects of these factors may differ depending on whether the gene is an oncogene versus a tumor suppressor gene. The chance that a variant is a driver may also depend on the flanking nucleotides, as represented by its single base substitution signature. It is our expectation that these factors will strongly influence the chance that a given variant is a driver. Our proposed method, by aggregating the evidence across all mutations in the data with similar such characteristics, will provide the evidence basis to make more confident inferences about the driver status of individual variants. In constructing this expanded modeling structure, we create a set of discrete variant sub-types that are defined by combinations of the covariates that we elect to include. This sub-type is denoted by the subscript "𝑙" in the various formulas below and is nested within a gene *j*. By doing so the number of parameters will be expanded substantially. To accommodate this additional complexity, we adopt a Bayesian hierarchical modeling framework, which offers flexible model specification and effectively handles complex structures without overfitting. We refer to our new method as BayesMAGPIE. A key advantage of using Bayesian hierarchical modeling is its ability to control, using prior distributions, the sparsity of nominated drivers, especially in settings with limited sample sizes. This approach is motivated by the general principle that each tumor type is typically driven by a relatively small number of genes within a pathway.

The complete data generation algorithm which defines this model is as follows.

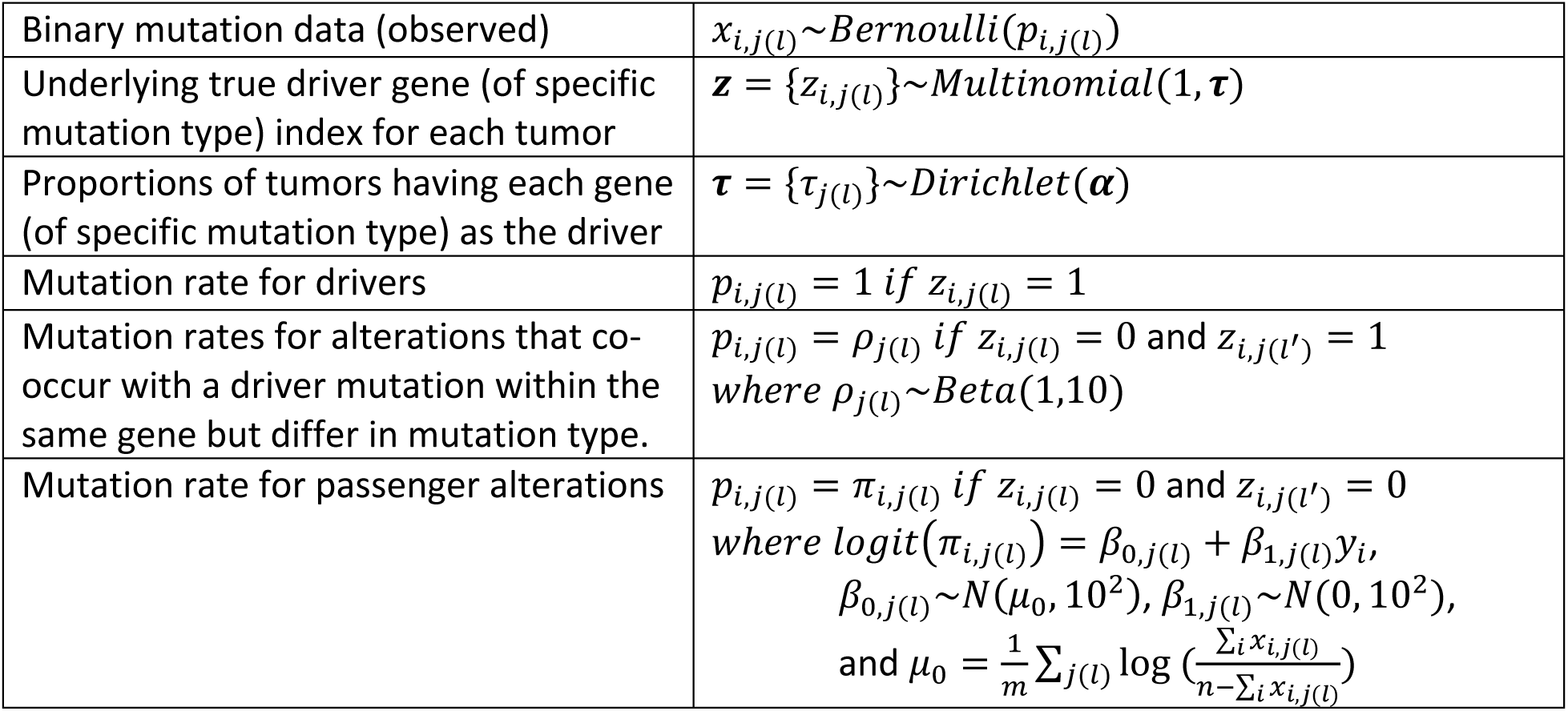

|  |  |
| --- | --- |
| Binary mutation data (observed) | $x_{i,j(l)} \sim \text{Bernoulli}(p_{i,j(l)})$ |
| Underlying true driver gene (of specific mutation type) index for each tumor | $\mathbf{z} = \{z_{i,j(l)}\} \sim \text{Multinomial}(1, \boldsymbol{\tau})$ |
| Proportions of tumors having each gene (of specific mutation type) as the driver | $\boldsymbol{\tau} = \{\tau_{j(l)}\} \sim \text{Dirichlet}(\boldsymbol{\alpha})$ |
| Mutation rate for drivers | $p_{i,j(l)} = 1 \text{ if } z_{i,j(l)} = 1$ |
| Mutation rates for alterations that co-occur with a driver mutation within the same gene but differ in mutation type. | $p_{i,j(l)} = \rho_{j(l)} \text{ if } z_{i,j(l)} = 0 \text{ and } z_{i,j(l')} = 1$<br>where $\rho_{j(l)} \sim \text{Beta}(1, 10)$ |
| Mutation rate for passenger alterations | $p_{i,j(l)} = \pi_{i,j(l)} \text{ if } z_{i,j(l)} = 0 \text{ and } z_{i,j(l')} = 0$<br>where $\text{logit}(\pi_{i,j(l)}) = \beta_{0,j(l)} + \beta_{1,j(l)} y_i$,<br>$\beta_{0,j(l)} \sim N(\mu_0, 10^2)$, $\beta_{1,j(l)} \sim N(0, 10^2)$,<br>and $\mu_0 = \frac{1}{m} \sum_{j(l)} \log \left( \frac{\sum_i x_{i,j(l)}}{n - \sum_i x_{i,j(l)}} \right)$ |

The mutual exclusivity of driver mutations is assumed only between different genes but not between mutation types within the same gene. This avoids falsely nominating driver genes whose missense and non-missense mutations form artificial perfect or near-perfect mutual exclusivity patterns. In other words, the passenger mutation rate of a gene is estimated based on its co-occurrence patterns with other genes, while the co-occurrence of different mutation types within the same gene is modeled using an additional set of nuisance parameters (𝜌_*j*(𝑙)_), which represent the conditional probability of observing a mutation of type 𝑙 in gene *j*, given that the tumor already harbors a driver mutation in the same gene but of a different type 𝑙^′^. Since the number of such co-occurring mutations is typically extremely low within a gene, these parameters 𝜌_*j*(𝑙)_ may not be estimated well with finite samples. This would make it difficult to determine which specific variant is the true driver when a tumor harbors both a missense and a non-missense mutation in the same gene. In practice, we recommend flagging such tumors and remaining cautious when interpreting which mutation type (e.g., missense vs. truncating) is the true driver in those tumors. Importantly, our method remains valid for identifying that a driver event exists in the gene in such tumors, even if its specific mutation type is undecided.

Moreover, because such co-occurring mutations within a gene are relatively rare, the estimation of 𝜌_*j*(𝑙)_ has minimal impact on the overall driver frequency estimates for each gene. By default, hyperparameters are chosen to define weakly informative prior distributions: 𝜌_*j*(𝑙)_∼𝐵𝑒𝑡𝑎(1,10), 𝛽_0,*j*(𝑙)_∼𝑁(𝜇_0_, 10^2^), and 𝛽_1,*j*(𝑙)_∼𝑁(0, 10^2^). We set 𝜇_0_ = 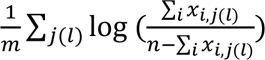, a crude average of the log-odds of the observed relative mutation frequencies across all genes. Here, *n* is the total number of tumors, and *m* is the total number of features, where each feature is defined as either a gene not categorized by mutation type or a specific mutation type within a gene. The hyperparameter set 𝜶 (i.e., 𝝉 = {𝜏_*j*(𝑙)_}∼𝐷*i*𝑟*i*𝑐ℎ𝑙𝑒𝑡(𝜶)) governs the prior distribution over the driver probabilities for each gene (of certain mutation type). Since we do not incorporate prior biological knowledge on driver probabilities, we adopt a uniform prior by assigning the same weight to each component (i.e., 𝛼_*j*(𝑙)_ = 𝛼_0_ for all *j*(𝑙)). The choice of 𝛼_0_ also influences the sparsity and concentration of the driver probabilities as follows:

- Larger values (e.g., 𝛼_0_ > 1) impose more informative prior beliefs, concentrating the posterior around the prior mean (i.e., 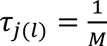, which can obscure true driver patterns.
- Smaller values (e.g., 𝛼_0_ < 0.1) yield less informative priors, allowing the data to drive a sparser and more selective driver nomination.

We will evaluate the impact of different choices for the hyperparameter 𝛼_0_on driver identification through both simulation studies and data applications to provide practical guidance for users. In general, we will demonstrate smaller values (e.g., 𝛼_0_ = 0.1, 0.01 or 0.001) tend to yield better overall performance.

To address the computational challenges associated with MCMC in the Bayesian hierarchical modeling framework, we will use variational inference(Hoffman, Blei, Wang and Paisley, 2013) for posterior estimation. Unlike MCMC, which relies on iteratively generating samples that asymptotically converge to the true posterior distribution, variational inference reformulates posterior inference as an optimization problem, typically resulting in significantly faster convergence. Variational inference seeks the best approximation to the true posterior within a chosen family of distributions by minimizing the Kullback-Leibler divergence from the variational distribution to the true posterior. For numerical optimization, we use the Adaptive Moment Estimation (Adam) algorithm(Kingma and Ba, 2014), a stochastic optimization method widely used in deep learning and machine learning. From the variational approximation, we will obtain the maximum a posteriori estimates of each model parameter (e.g., 𝜏_*j*(𝑙)_, 𝜌_*j*(𝑙)_, 𝛽_0,*j*(𝑙)_ and 𝛽_1,*j*(𝑙)_). Using these estimates, we will further predict the most likely driver event (or none) for each tumor by computing the posterior probability, 𝑝(𝑧_*ij*(𝑙)_ = 1|*x*_*i*_, 𝑦_*i*_), conditional on the observed mutation profile and relevant covariates (e.g., TMB).

In the following sections, we first illustrate our proposed method using real datasets, followed by simulation studies to demonstrate its operating characteristics.

## 3 DATA APPLICATIONS

We apply our method to The Cancer Genome Atlas Lung Adenocarcinoma (TCGA-LUAD) dataset and conduct an analysis on the RTK-RAS pathway to illustrate how BayesMAGPIE works and what it delivers. We then extend our analysis to seven additional cancer types from TCGA, benchmarking BayesMAGPIE’s performance against the original MAGPIE.

### 3.1 Illustration of BayesMAGPIE: Analyzing the RTK-RAS Pathway in TCGA-LUAD

#### 3.1.1 Data Preparation

For our illustration of the method, we focus solely on mutations in the RTK-RAS pathway, a major known pathway that influences the development of LUAD. Pathway genes are defined based on the framework established by Sanchez-Vega et al (2018). This study cohort includes 502 tumors and 37 RTK-RAS pathway genes. To incorporate mutation variant types, we classify non-synonymous mutations in each gene within each tumor as either missense or other (e.g., nonsense mutations, frameshift deletions) and construct a binary tumor-by-gene-variant mutation matrix. Notably, this classification is not mutually exclusive, meaning a tumor can harbor both mutation types within the same gene. Since missense mutations are often predominant, we introduce a prescreening rule to balance feature dimensionality and computational stability. Specifically, we classify genes by variant type only if both missense and other mutations occur in at least min(10, 0.01*n*) tumors, where *n* is the total number of tumors. After applying this criterion, the final dataset (denoted by 𝑋_𝑣𝑎𝑟*i*𝑎*n*𝑡_) comprises 47 features: 27 genes without variant type classification and 10 genes distinguished by missense/other classification. We also compare our results with two additional scenarios: (1) analysis using MAGPIE and (2) analysis using BayesMAGPIE without incorporating variant type information. In both cases, we use gene-level binary mutation data (37 features), denoted as^𝑋^𝑔𝑒*n*𝑒.

#### 3.1.2 Analysis Plan

We conduct three sets of analyses: 1) MAGPIE on 𝑋_𝑔𝑒*n*𝑒_; 2) BayesMAGPIE on 𝑋_𝑔𝑒*n*𝑒_; and 3) BayesMAGPIE on 𝑋_𝑣𝑎𝑟*i*𝑎*n*𝑡_. For each BayesMAGPIE analysis, we further perform three sub-analyses, varying the hyperparameter 𝜶 in the Dirichlet distribution that models the relative frequency of driver mutations for each gene *j* (or each specific mutation type 𝑙 within gene *j*), i.e., 𝝉 = {𝜏_*j*(𝑙)_}∼𝐷*i*𝑟*i*𝑐ℎ𝑙𝑒𝑡(𝜶), 𝑤ℎ𝑒𝑟𝑒 𝜶 = {𝛼_0_, … , 𝛼_0_}. Specifically, we consider the following settings: 𝛼_0_=1, 𝛼_0_=0.1, 𝛼_0_=0.01, and 𝛼_0_=0.001, where the smaller density values (i.e., less informative prior) would yield sparser results. Both MAGPIE and BayesMAGPIE compute tumor-specific posterior probability vectors, summarizing the estimated relative frequency of driver mutations within each gene (of specific mutation type, when applicable) and the probability of harboring no driver mutation. These probabilities, along with the observed mutation relative frequency, will be visualized using waterfall plots and are summarized in tables for interpretation.

#### 3.1.3 Results

The data reveal that 83% of the 502 tumors had a mutation in the RTK-RAS pathway in TCGA-LUAD. In MAGPIE analysis, we estimated that in 65% of the tumors one of the mutations is considered to be the driver. Figure 1A displays data (top panel) and model estimates (middle panel) for the nine genes with an estimated values of 𝜏_*j*_>0.01 (at least 1% of tumors estimated to carry a driver mutation in gene *j*), along with six genes estimated to be non-driver (𝜏_*j*_<0.001) but with relatively high observed mutation relative frequency (>3.5%) as compared to other non-drivers. The reason we include additional non-driver genes is to demonstrate their noisier mutational patterns (more non-singletons) compared to estimated driver genes, where singletons refer to tumors harboring mutation in only one gene in a pathway. In the bottom panel we show the distribution of TMB (high vs. low), where we generally observe that a tumor with higher TMB is associated with less chance of harboring a driver mutation. The nine genes with relatively high driver probabilities are *KRAS*, *EGFR*, *NF1*, *BRAF*, *MET*, *ERBB3*, *RIT1*, *MAP2K1* and *SOS1*, among which *KRAS* and *EGFR* are well-known drivers in LUAD. The observed mutation relative frequencies and estimated driver relative frequencies for each gene are summarized in Table 1A. Figure 1B and 1C display the result for the analysis with BayesMAGPIE under the setting of not incorporating mutation type but with two different Dirichlet parameter values, 𝛼_0_ = 1 and 𝛼_0_ = 0.1, respectively. With a more informative prior, i.e., 𝛼_0_ = 1, the result is similar to MAGPIE, while with a less informative prior, i.e., 𝛼_0_ = 0.1, the result becomes sparser leading to a smaller number of identified driver genes (5 genes: *KRAS*, *EGFR*, *NF1*, *BRAF*, *MET*). We also observe that the result generally remains unchanged with even less informative priors, e.g., 𝛼_0_ = 0.01 and 𝛼_0_ = 0.001, as shown in Table 1A. Thus, we demonstrate that judicious selection of the degree of informativeness of the prior in BayesMAGPIE has the effect of limiting the number of genes in a pathway to be called as drivers.

**FIGURE 1.**
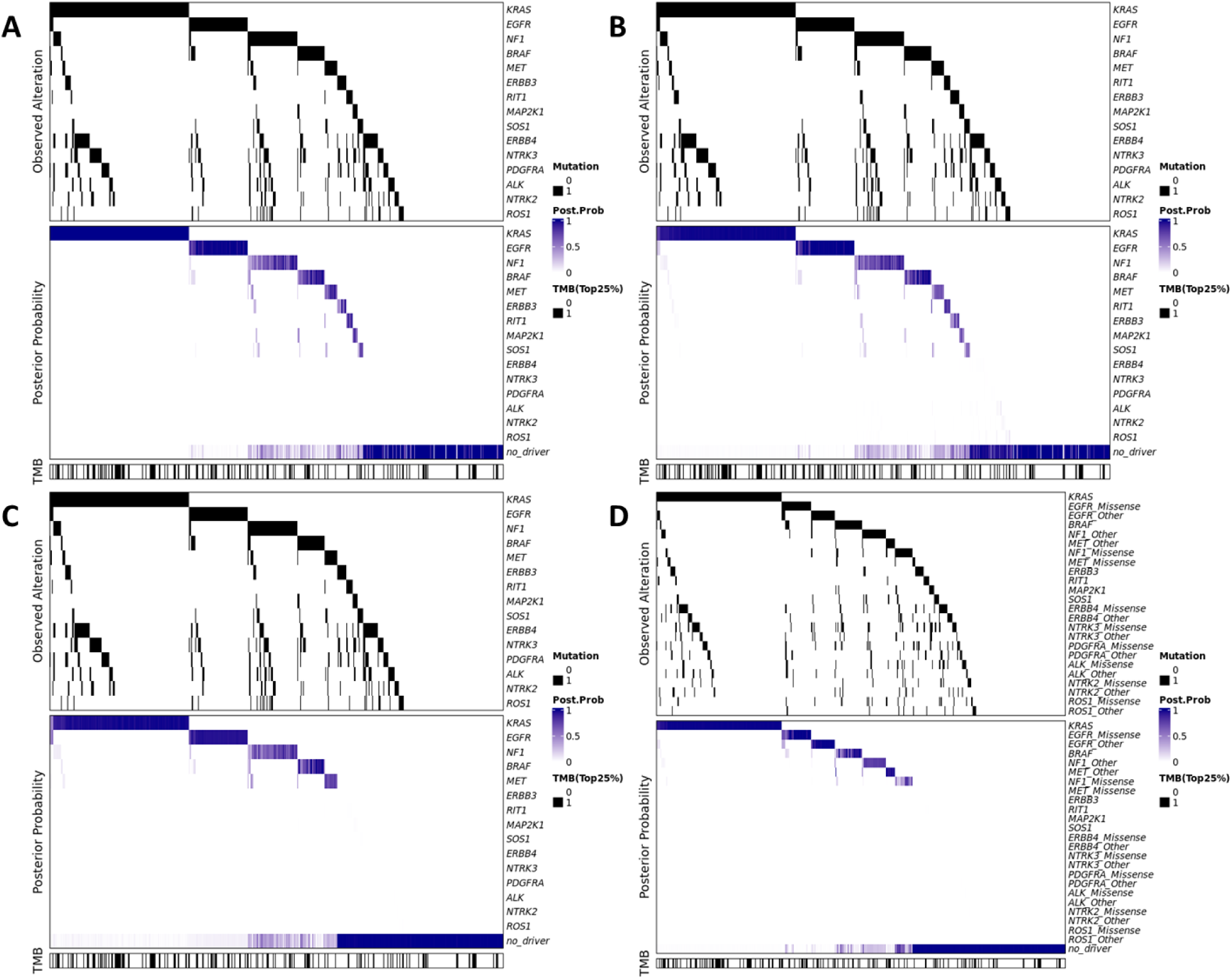
Illustration of the Observed Binary Mutation Status, Estimated Posterior Probability of Driver Mutation, and the Distribution of Binary TMB of RTK-RAS Pathway in LUAD (Showing a Subset of Genes). A. MAGPIE; B. BayesMAGPIE not incorporating mutation type (𝛼_0_=1); C. BayesMAGPIE not incorporating mutation type (𝛼_0_=0.1); D. BayesMAGPIE incorporating mutation type (𝛼_0_=0.1).

**TABLE 1.** Summary of the Observed Mutation Frequency and the Estimated Driver Frequency for Each Gene in RTK-RAS Pathway (Top Genes Only) from TCGA-LUAD.

A. MAGPIE and BayesMAGPIE (Not Incorporating Mutation Type)
| Gene | Observed Mutation Freq | Inferred Driver Freq |  |  |  |  |
| --- | --- | --- | --- | --- | --- | --- |
| | | MAGPIE | BayesMAGPIE $\alpha_0=1$ | BayesMAGPIE $\alpha_0=0.1$ | BayesMAGPIE $\alpha_0=0.01$ | BayesMAGPIE $\alpha_0=0.001$ |
| *KRAS* | 0.307 | 0.307 | 0.300 | 0.293 | 0.295 | 0.295 |
| *EGFR* | 0.137 | 0.124 | 0.126 | 0.125 | 0.128 | 0.128 |
| *NF1* | 0.129 | 0.072 | 0.078 | 0.073 | 0.073 | 0.073 |
| *BRAF* | 0.084 | 0.050 | 0.053 | 0.050 | 0.051 | 0.051 |
| *MET* | 0.046 | 0.023 | 0.019 | 0.025 | 0.025 | 0.025 |
| *ERBB3* | 0.038 | 0.013 | 0.012 | <0.001 | <0.001 | <0.001 |
| *RIT1* | 0.022 | 0.011 | 0.012 | <0.001 | <0.001 | <0.001 |
| *MAP2K1* | 0.018 | 0.010 | 0.008 | <0.001 | <0.001 | <0.001 |
| *SOS1* | 0.032 | 0.010 | 0.008 | <0.001 | <0.001 | <0.001 |

B. BayesMAGPIE Incorporating Mutation Type
|  |  | Inferred Driver Freq |  |  |  |
| --- | --- | --- | --- | --- | --- |
| Gene | Observed Mutation Freq | BayesMAGPIE $\alpha_0=1$ | BayesMAGPIE $\alpha_0=0.1$ | BayesMAGPIE $\alpha_0=0.01$ | BayesMAGPIE $\alpha_0=0.001$ |
| *KRAS* | 0.307 | 0.303 | 0.303 | 0.303 | 0.303 |
| *EGFR* |  |  |  |  |  |
| Missense | 0.074 | 0.067 | 0.065 | 0.065 | 0.065 |
| Other | 0.072 | 0.039 | 0.059 | 0.059 | 0.059 |
| *NF1* |  |  |  |  |  |
| Missense | 0.058 | 0.020 | 0.020 | 0.020 | 0.020 |
| Other | 0.076 | 0.042 | 0.045 | 0.046 | 0.046 |
| *BRAF* | 0.084 | 0.053 | 0.051 | 0.051 | 0.051 |
| *MET* |  |  |  |  |  |
| Missense | 0.020 | <0.001 | <0.001 | <0.001 | <0.001 |
| Other | 0.026 | 0.021 | 0.024 | 0.024 | 0.024 |
| *ERBB3* | 0.038 | 0.011 | <0.001 | <0.001 | <0.001 |
| *RIT1* | 0.022 | 0.010 | <0.001 | <0.001 | <0.001 |
| *MAP2K1* | 0.018 | 0.005 | <0.001 | <0.001 | <0.001 |
| *SOS1* | 0.032 | 0.006 | <0.001 | <0.001 | <0.001 |

For the analysis incorporating mutation types (missense vs. other), the results are presented in Figure 1D and Table 1B. We observe that driver mutations in *EGFR* are slightly more enriched in missense mutations (likely due to the hotspot driver variant L858R), whereas driver mutations in *NF1* show a moderate enrichment in non-missense mutation types. Notably, nearly all driver mutations in *MET* are estimated to be of types other than missense. To further investigate, we examine individual *MET* variants and summarize the average probability of harboring a driver mutation in *MET* in tumors where the corresponding variant is observed, as an indicator of its potential importance (Table 2). Among the top-ranked variants identified by BayesMAGPIE are the splice site mutations, including p.X879_splice, p.X1028_splice, and p.X1027_splice. Notably, p.X1028_splice and p.X1027_splice mutations are associated with *MET* exon 14 skipping, a clinically actionable event targeted by *MET* inhibitors such as Capmatinib and Tepotinib in non-small cell lung cancer (Paik et al., 2020; Wolf et al., 2020). While these two splice variants are also ranked high by MAGPIE, the average estimated driver probability for p.X1028_splice is substantially lower compared to that from BayesMAGPIE. However, a more notable difference is the consistently low driver probabilities from BayesMAGPIE for the missense mutations.

**TABLE 2.** Variant Analysis – *MET* in TCGA-LUAD.

| MAGPIE | | | | BayesMAGPIE<br>(Mutation type incorporated, $\alpha_0=0.1$) | | | |
| --- | --- | --- | --- | --- | --- | --- | --- |
| Variant | Type | Freq | Prob | Variant | Type | Freq | Prob |
| p.Y1021* | Nonsense | 1 | 0.955 | p.X879_splice | Splice_Site | 1 | 0.979 |
| p.X1027_splice | Splice_Site | 1 | 0.943 | p.X1028_splice | Splice_Site | 6 | 0.949 |
| p.E1332G | Missense | 1 | 0.857 | p.X1027_splice | Splice_Site | 1 | 0.926 |
| p.X1028_splice | Splice_Site | 6 | 0.776 | p.Y1021* | Nonsense | 1 | 0.915 |
| p.S441C | Missense | 1 | 0.748 | . | Frame_Shift_Del | 1 | 0.914 |
| . | Frame_Shift_Del | 1 | 0.471 | p.S441C | Missense | 1 | 0.002 |
| p.L317I | Missense | 1 | 0.469 | p.E1332G | Missense | 1 | 0.002 |
| p.Y1253H | Missense | 1 | 0.424 | p.L317I | Missense | 1 | 0.001 |
| p.N315S | Missense | 1 | 0.263 | p.N315S | Missense | 1 | 0.001 |
| p.R412H | Missense | 1 | 0.23 | p.R412H | Missense | 1 | 0.001 |
| p.X879_splice | Splice_Site | 1 | 0.2 | p.Y1253H | Missense | 1 | 0.001 |
| p.C98F | Missense | 1 | <0.001 | p.H476Y | Missense | 1 | 0.001 |
| p.T660R | Missense | 1 | <0.001 | p.D208Y | Missense | 1 | 0.001 |
| p.D208Y | Missense | 1 | <0.001 | p.T660R | Missense | 1 | <0.001 |
| p.H476Y | Missense | 1 | <0.001 | p.C98F | Missense | 1 | <0.001 |

### 3.2 Cross-Cancer Evaluation of BayesMAGPIE Using TCGA Data

To further evaluate BayesMAGPIE’s performance in real-world settings, we applied it to a comprehensive set of independent analyses of the RTK-RAS pathway across eight cancer types using TCGA data. Both MAGPIE and BayesMAGPIE were analyzed as before. For BayesMAGPIE, we used binary gene alteration data that incorporated mutation types (missense vs. other) and examined two prior settings to assess the impact of hyperparameter choices: a relatively more informative Dirichlet prior (𝛼_0_ = 1), and a less informative prior (𝛼_0_= 0.1). This comparison highlights how the prior influences the resulting driver mutation estimates.

Table 3 summarizes key descriptive statistics and model estimates for each tumor type, including the number of tumors analyzed (# of Tumors), the relative frequency of observed mutations (Mut Freq), and the estimated proportion of tumors harboring a driver mutation in the RTK-RAS pathway (Inferred Driver Freq). In addition to lung adenocarcinoma (LUAD), which was discussed earlier, the MAGPIE results highlight that cutaneous melanoma (SKCM) and papillary thyroid carcinoma (THCA) also exhibit high frequencies of driver mutations in the RTK-RAS pathway. These are followed by urothelial bladder cancer (BLCA) and uterine corpus endometrial carcinoma (UCEC), where around 40% of tumors are estimated to harbor a driver mutation in the pathway. In contrast, breast cancer (BRCA), lower grade glioma (LGG), and head and neck squamous cell carcinoma (HNSC) show relatively low proportions of RTK-RAS drivers. The results from BayesMAGPIE are similar to those of MAGPIE when using a more informative prior (𝛼_0_ = 1). However, selecting a less informative prior (𝛼_0_= 0.1) yields a sparser set of nominated drivers, in which BRCA, LGG, and HNSC are estimated to have extremely low driver relative frequencies. This suggests that mutations in the RTK-RAS pathway are less likely to act as drivers in BRCA, LGG, and HNSC, where other pathways may play a more prominent role, compared to other tumor types such as LUAD, SKCM, and THCA.

**TABLE 3.**
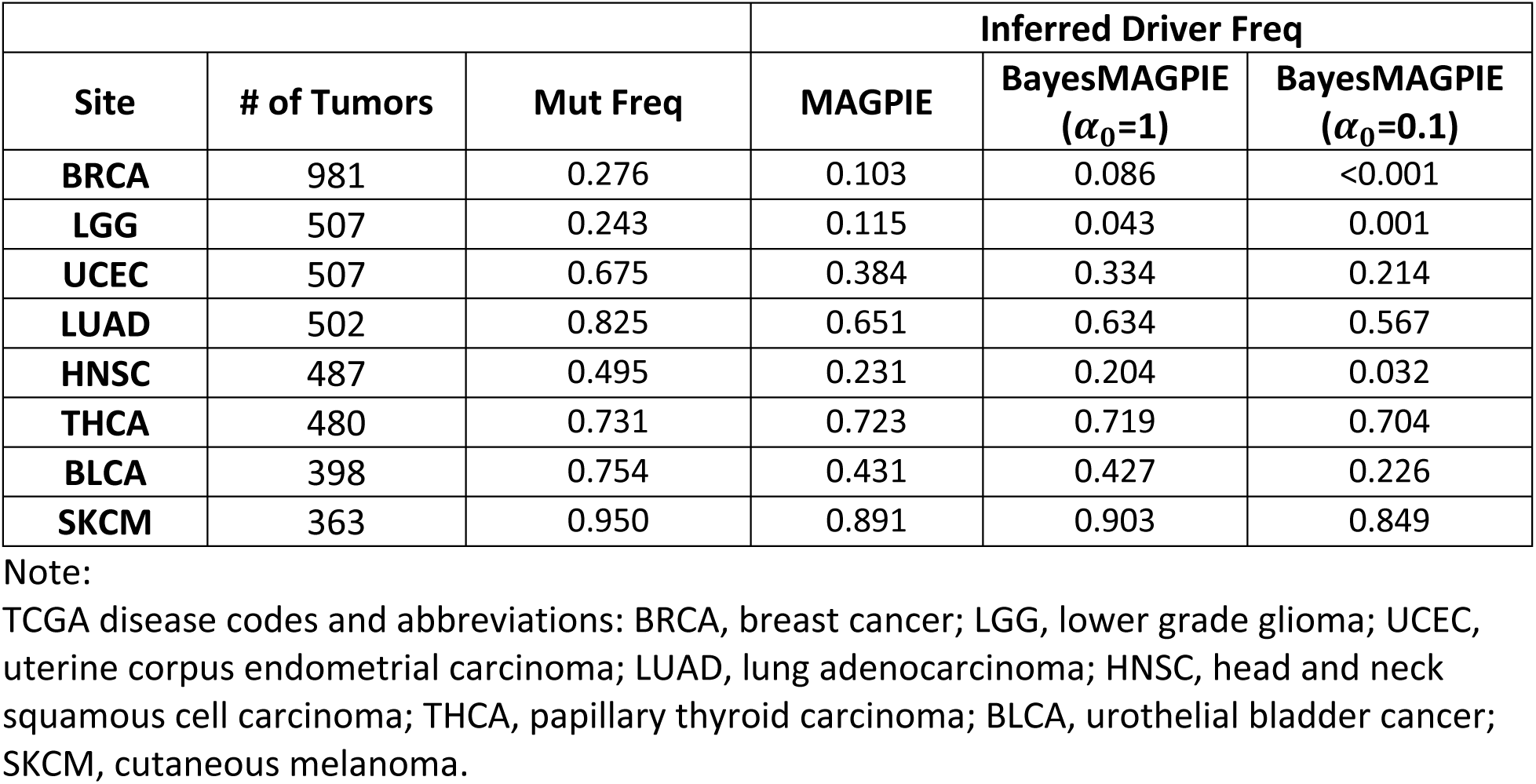
Results of Independent Analyses on RTK-RAS Pathway in Each Tumor Site.

| Site | # of Tumors | Mut Freq | Inferred Driver Freq |  |  |
| --- | --- | --- | --- | --- | --- |
| | | | MAGPIE | BayesMAGPIE ($\alpha_0=1$) | BayesMAGPIE ($\alpha_0=0.1$) |
| BRCA | 981 | 0.276 | 0.103 | 0.086 | <0.001 |
| LGG | 507 | 0.243 | 0.115 | 0.043 | 0.001 |
| UCEC | 507 | 0.675 | 0.384 | 0.334 | 0.214 |
| LUAD | 502 | 0.825 | 0.651 | 0.634 | 0.567 |
| HNSC | 487 | 0.495 | 0.231 | 0.204 | 0.032 |
| THCA | 480 | 0.731 | 0.723 | 0.719 | 0.704 |
| BLCA | 398 | 0.754 | 0.431 | 0.427 | 0.226 |
| SKCM | 363 | 0.950 | 0.891 | 0.903 | 0.849 |

Note:
TCGA disease codes and abbreviations: BRCA, breast cancer; LGG, lower grade glioma; UCEC, uterine corpus endometrial carcinoma; LUAD, lung adenocarcinoma; HNSC, head and neck squamous cell carcinoma; THCA, papillary thyroid carcinoma; BLCA, urothelial bladder cancer; SKCM, cutaneous melanoma.

Following the overall pathway-level driver illustration, Figure 2 provides a detailed overview of the specific driver genes identified in each tumor site. In the graph, the size of each bubble reflects the relative frequency of mutations associated with a specific gene (y-axis) within the corresponding cancer type (x-axis). The depth of the bubble’s color indicates the ratio of the estimated driver mutation rate to the observed mutation rate for that gene, effectively representing the conditional probability that an observed mutation is classified as a driver alteration. Strong driver genes are expected to exhibit a darker shade, allowing for easy visual recognition. The three subpanels correspond to results from the three analytical approaches introduced earlier: A, MAGPIE; B, BayesMAGPIE with a more informative prior (𝛼_0_ = 1); and C, BayesMAGPIE with a less informative prior (𝛼_0_= 0.1). Consistent with earlier observations, we find that Fig. 2A and 2B display noisier estimated driver patterns, while BayesMAGPIE with a less informative prior produces a notably sparser set of results. For example, in BLCA MAGPIE estimates more than ten genes as drivers in a high proportion of tumors harboring mutations in them, whereas BayesMAGPIE with a less informative prior highlights only four prominent drivers: *HRAS*, *KRAS*, *FGFR3*, and *FGFR2*. Across other tumor types, the sparse list of key drivers identified by BayesMAGPIE includes: *NRAS* in SKCM, THCA and UCEC (weak driver); *NF1* in LUAD and SKCM (weak driver); *MET* in LUAD; *KRAS* in BLCA, LUAD and UCEC (weak driver); *KIT* in SKCM; *HRAS* in BLCA and THCA; *FGFR3* in BLCA; *FGFR2* in BLCA and UCEC (weak driver); *EGFR* in LUAD; *BRAF* in LUAD, SKCM and THCA; and *ALK* in HNSC.

**FIGURE 2.**
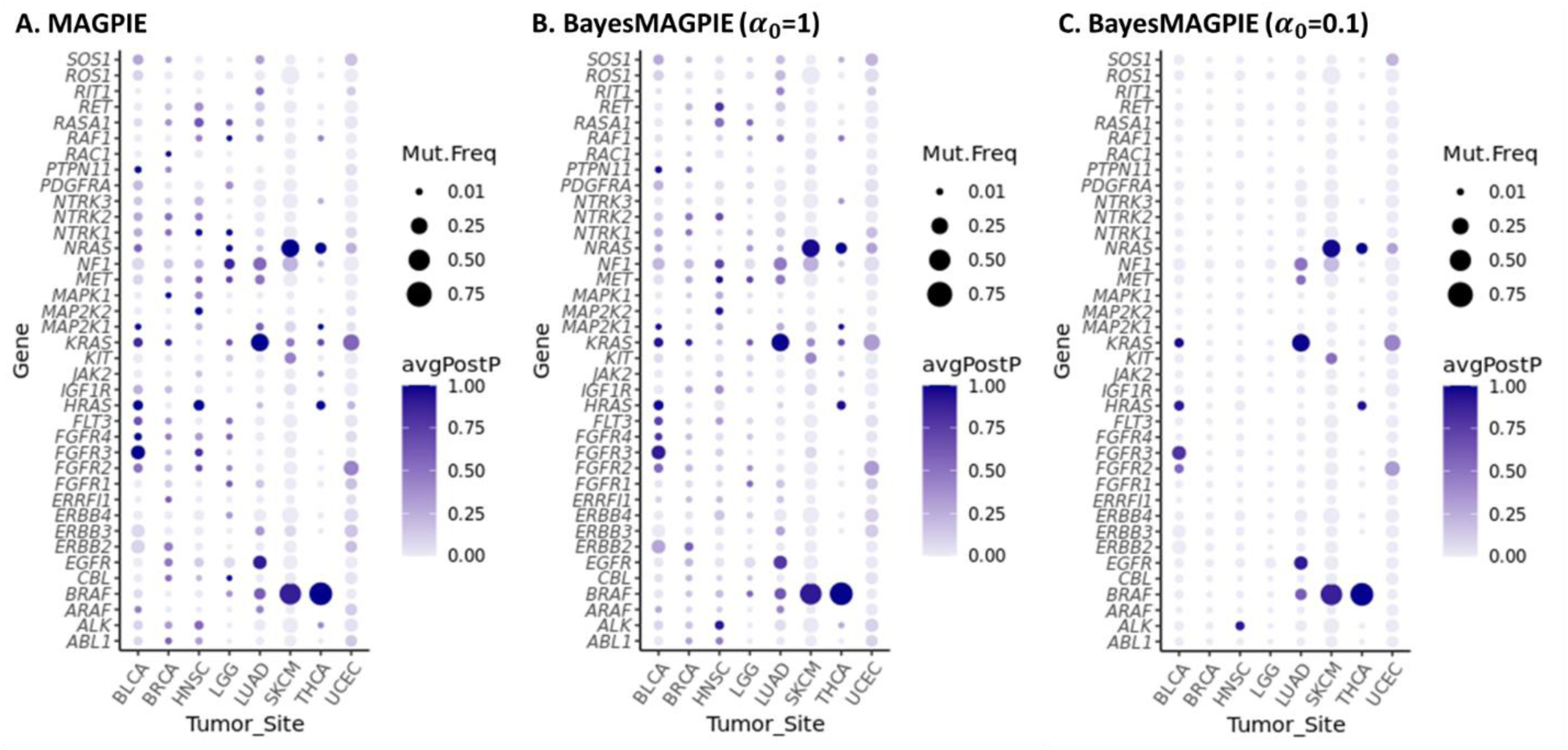
Bubble Plots Showing the Likelihood of Individual Genes within RTK-RAS Pathway as Driver across Different Tumor Sites. A: Result from MAGPIE; B: Result from BayesMAGPIE with a more informative prior; C: Result from BayesMAGPIE with a less informative prior.

We believe that the capability of BayesMAGPIE to control the sparsity of driver nomination through the prior distribution is a significant advantage in real-world applications, particularly when researchers have limited resources to experimentally validate novel driver genes. By prioritizing a smaller, more selective list of candidates, BayesMAGPIE increases the likelihood that nominated drivers are truly functional. The operating characteristics of BayesMAGPIE, including bias in estimation, false positive rates, and true positive rates, are evaluated in the following section through simulation studies.

## 4 OPERATING CHARACTERISTICS OF THE METHOD

### 4.1 Simulation Settings

We conducted simulations to further evaluate the performance of the proposed method. Data were generated based on the aforementioned algorithm (Proposed Approach), with parameters including 𝝉, 𝜷_𝟎_, 𝜷_𝟏_ and 𝝆 estimated from TCGA-LUAD data (RTK-RAS pathway) to reflect real-world scenarios. To simplify the simulation setup, we modeled 10 genes in total, consisting of 5 drivers and 5 non-drivers. Details of their overall and driver relative frequencies (i.e., 𝝉) are provided in Table 4. The covariate TMB was simulated from a standard normal distribution to mirror the centered log-scaled scores used in the model. We varied the number of tumors from 200 to 2000 to reflect typical sample sizes and varied the Dirichlet hyperparameter (𝛼_0_) from 1 to 0.001 to assess the impact of the prior distribution on driver detection. For each setting, 500 data replicates were generated under the proposed model structure.

**TABLE 4.** Characteristics of Simulated Genes.

|  | Simulated Gene Index<br>(Variant Type) | Real Gene<br>Represented | Simulated<br>Mutation Freq | Simulated<br>Driver Freq |
| --- | --- | --- | --- | --- |
| Driver | Gene 0 | *KRAS* | 0.306 | 0.302 |
|  | Gene 1 | *BRAF* | 0.084 | 0.050 |
|  | Gene 2 | *EGFR* |  |  |
|  | Missense |  | 0.076 | 0.069 |
|  | Other |  | 0.083 | 0.032 |
|  | Gene 3 | *NF1* |  |  |
|  | Missense |  | 0.056 | 0.020 |
|  | Other |  | 0.078 | 0.035 |
|  | Gene 4 | *MET* |  |  |
|  | Missense |  | 0.020 | 0 |
|  | Other |  | 0.025 | 0.025 |
| Non-Driver | Gene 5 | *ERBB4* |  |  |
|  | Missense |  | 0.072 | 0 |
|  | Other |  | 0.042 | 0 |
|  | Gene 6 | *NTRK3* |  |  |
|  | Missense |  | 0.081 | 0 |
|  | Other |  | 0.035 | 0 |
|  | Gene 7 | *PDGFRA* |  |  |
|  | Missense |  | 0.064 | 0 |
|  | Other |  | 0.038 | 0 |
|  | Gene 8 | *ALK* |  |  |
|  | Missense |  | 0.049 | 0 |
|  | Other |  | 0.040 | 0 |
|  | Gene 9 | *NTRK2* |  |  |
|  | Missense |  | 0.029 | 0 |
|  | Other |  | 0.033 | 0 |

### 4.2 Bias in Estimated Driver Frequencies

We first evaluated the bias of the maximum a posteriori estimates of our core model parameters, i.e., the relative driver frequency for each gene (of a specific mutation type), denoted as 𝝉 = {𝜏_*j*(𝑙)_}. The true 𝜏_*j*(𝑙)_ values are summarized in Table 4 ("Simulated Driver Frequency"). In Figure 3, we observe that when 𝛼_0_= 1 (a more informative prior), the non-driver genes (genes 5-9) tend to consistently show a small positive bias (i.e., 𝜏^_*j*(𝑙)_>0) with smaller sample sizes (e.g., *n* = 200 and 500), likely leading to false positive findings (see 4.3). The detailed bias values are provided in Supplemental Table 1. This pattern is consistent with observations from previous real data applications, where a larger number of genes were nominated as drivers under more informative prior settings. Such bias among non-drivers is substantially reduced when a less informative prior is used (𝛼_0_=0.1) even with a small sample size, though this comes at the cost of increased negative bias among true driver genes (genes 0-4). For example, gene 1, whose true driver frequency is 0.05, shows an average bias shift from 0 to -0.006 as the prior becomes less informative (*n* = 200, Supplemental Table 1A-B). In other words, the sensitivity for detecting true drivers is likely to decrease (see 4.3). We also observe that the performance of BayesMAGPIE stabilizes as 𝛼_0_decreases below 0.1 (Supplemental Fig.1B-D, Supplemental Table 1B-D). Therefore, in practice, setting 𝛼_0_= 0.1 offers a reasonable sparse solution. Lastly, as the sample size increases, the impact of the prior diminishes. Based on our simulations, with 500 to 1000 tumors, BayesMAGPIE provides reasonably accurate estimates of relative driver frequencies with a less informative prior.

**FIGURE 3.**
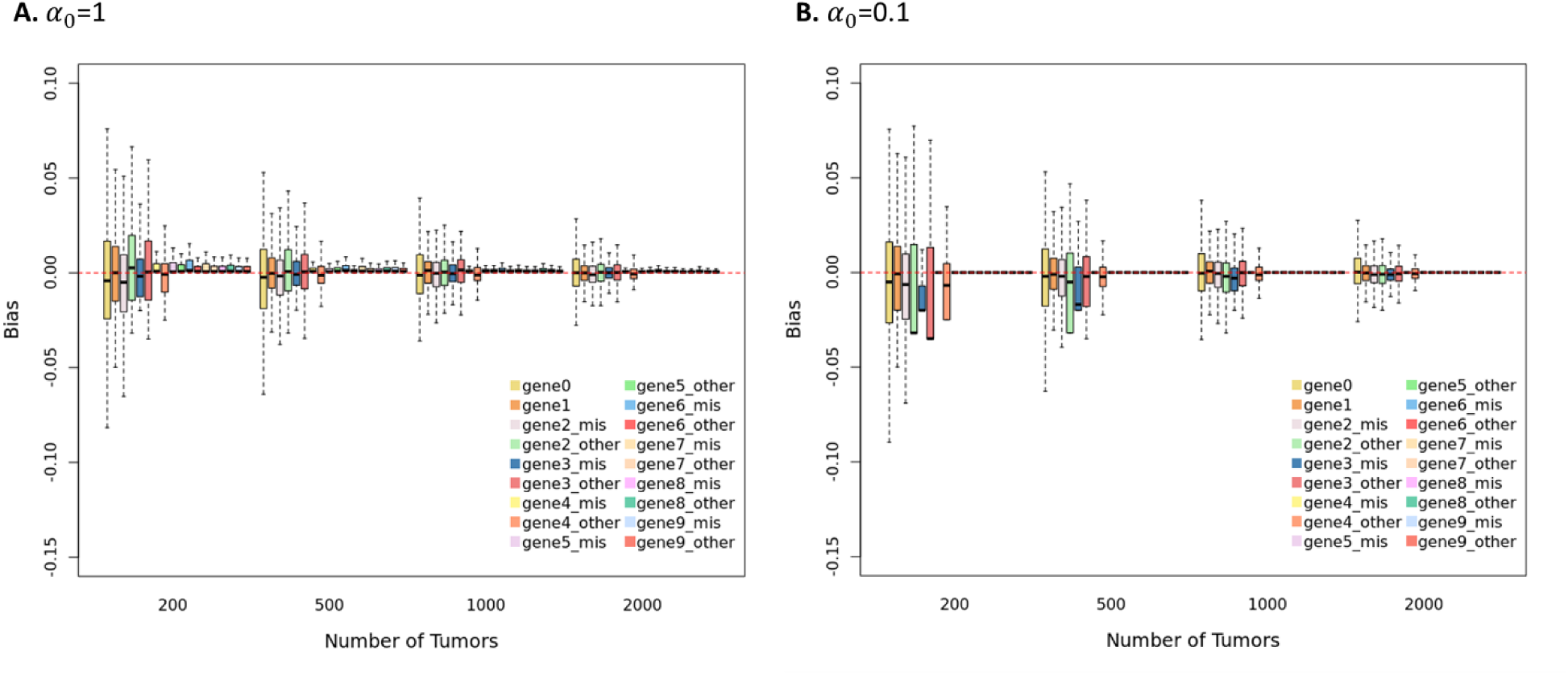
Boxplots Showing the Bias in the Estimated Driver Frequency of Each Gene from BayesMAGPIE under Varying Dirichlet Priors and Different Sample Sizes. A: 𝛼_0_=1; B: 𝛼_0_=0.1.

### 4.3 Driver Prediction Accuracy

Next, we evaluated the accuracy of the proposed method for identifying drivers in individual tumors. First, we assessed the overall false positive rate (FPR) for predicting a driver in tumors that do not harbor any driver mutation under our simulation design. The FPR is calculated as:

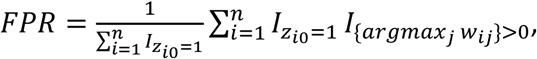

where 𝑧_*i*0_=1 indicates that tumor *i* does not harbor a driver mutation, and 𝑤_*ij*_ = 𝑝(𝑧_*ij*_ = 1|*x*_*i*_, 𝑦_*i*_) denotes the posterior probability that tumor *i* harbors a driver mutation in gene *j*, conditional on the observed mutation profile *x*_*i*_ and relevant covariate 𝑦_*i*_ (e.g., TMB).

Similarly, we evaluated the true positive rate (TPR) for correctly identifying the true driver gene among tumors that do harbor a driver mutation. The TPR is calculated as:

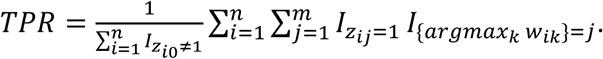

The results are summarized in Table 5, where the FPR and TPR are averaged across 500 simulated data replicates. With a large sample size (*n* = 2000), the influence of the prior is minimal, and the corresponding FPR (∼0.1) and TPR (∼0.95) reflect the optimal performance of the proposed method under our simulation design. Using this as a benchmark, we observe that with smaller sample sizes, employing a less informative prior reduces the FPR but at the cost of a lower TPR, which is consistent with the results observed for estimation bias in 4.2. For example, when *n* = 200, the FPR drops from 0.160 to 0.067 as the prior becomes less informative, while the TPR decreases from 0.921 to 0.830. However, this reduction in TPR is largely mitigated by increasing the sample size. With 500 to 1000 tumors, the TPR improves to approximately 0.90-0.93 while keeping the FPR below 0.1.

**TABLE 5.** Overall Accuracy of Driver Prediction.

| | Sample Size | Dirichlet Prior<br>$\alpha_0$ | Accuracy | |
| --- | --- | --- | --- | --- |
|  |  |  | FPR | TPR |
| **MAGPIE** | 200 | N/A | 0.162 | 0.921 |
|  | 500 |  | 0.144 | 0.939 |
|  | 1000 |  | 0.141 | 0.945 |
|  | 2000 |  | 0.138 | 0.948 |
| **BayesMAGPIE** | 200 | 1 | 0.160 | 0.921 |
|  |  | 0.1 | 0.071 | 0.841 |
|  |  | 0.01 | 0.067 | 0.832 |
|  |  | 0.001 | 0.067 | 0.830 |
|  | 500 | 1 | 0.129 | 0.941 |
|  |  | 0.1 | 0.083 | 0.903 |
|  |  | 0.01 | 0.081 | 0.897 |
|  |  | 0.001 | 0.081 | 0.896 |
|  | 1000 | 1 | 0.121 | 0.953 |
|  |  | 0.1 | 0.095 | 0.933 |
|  |  | 0.01 | 0.094 | 0.931 |
|  |  | 0.001 | 0.093 | 0.930 |
|  | 2000 | 1 | 0.114 | 0.957 |
|  |  | 0.1 | 0.102 | 0.949 |
|  |  | 0.01 | 0.101 | 0.948 |
|  |  | 0.001 | 0.101 | 0.948 |

We further benchmarked the performance of BayesMAGPIE against our previously developed approach, MAGPIE. While MAGPIE tends to achieve a higher TPR with small to moderate sample sizes (e.g., *n* = 200 or 500), BayesMAGPIE (with less informative prior settings) shows a substantial advantage in controlling the FPR. This reflects a major strength of BayesMAGPIE in that it promotes sparsity in driver nomination and prioritizes a more selective list of candidate drivers for downstream experimental validation. Even with a large sample size (*n* = 2000), where both MAGPIE and BayesMAGPIE (regardless of the prior used) achieve a TPR of approximately 0.95, the FPR remains consistently lower for BayesMAGPIE (∼0.10) compared to MAGPIE (∼0.14). This improvement is largely attributed to BayesMAGPIE’s enhanced modeling framework, which incorporates mutation type information. This added flexibility allows the proposed method to distinguish between potential differences in driver roles between variant types within the same gene, thereby improving the accuracy of driver identification.

## 5 DISCUSSION

Our goal in developing this methodology was to provide an alternative strategy for identifying potential driver mutations in tumors and assigning probabilities to candidate events, with an emphasis on incorporating variant types and achieving better control of false positives to enhance driver identification. This new strategy builds on the concept of mutual exclusivity, a widely used framework for studying driver events. Among the many groups that have explored this framework, we chose to build on the work of Hua et al. (2016) and Wang et al. (2024) whose models are grounded in well-established statistical principles.

A key innovation of our new approach is the use of Bayesian hierarchical modeling to account for the complex structure of the data, enabling the incorporation of mutation-type information. By explicitly modeling mutation types (e.g., missense vs. truncating), our method can differentiate the functional roles of mutations within the same gene based on their type. In our detailed analysis of TCGA-LUAD data, we find that *MET* missense mutations are less likely to be drivers compared to other mutation types. Notably, our method identifies splice site mutations as the top variants in *MET*, including p.X1028_splice and p.X1027_splice, which are associated with *MET* exon 14 skipping, a clinically actionable event targeted by *MET* inhibitors such as Capmatinib and Tepotinib in non-small cell lung cancer (Paik et al., 2020; Wolf et al., 2020). These results highlight the potential of our method to pinpoint functionally relevant and potentially pathogenic mutations within specific genes.

Another key innovation of our new method is the use of a Dirichlet prior to model gene-specific driver frequencies, which allows effective control over the sparsity of inferred driver patterns through the tuning of hyperparameter values. This emphasis on sparse output is guided by the widely accepted principle that each tumor type is typically driven by a relatively small number of genes. Moreover, the reduced set of nominated driver genes and variants identified by our method can serve as strong candidates for experimental validation using modern in vitro and in vivo approaches (Hodis et al., 2022). Through extensive simulation studies, we find that a less informative prior (e.g., α = 0.1) consistently achieves a good balance of true positive rate (TPR) versus false positive rate (FPR) in datasets of moderate sample size, compared with other options.

We emphasize that our strategy focuses on a single pathway and relies on the key assumption that there is at most one driver mutation within that pathway in any given tumor, an assumption that may not always hold. Nevertheless, this assumption does provide a coherent and practical framework for analyzing mutual exclusivity. In contrast, it is well recognized that most tumors harbor multiple driver mutations, typically from different pathways. While our current approach can be applied independently to different pathways to identify a more comprehensive set of drivers, an important direction for future research is to extend the model to enable simultaneous analysis across multiple pathways.

## FUNDING

This study was supported by the National Institutes of Health (P01 CA206980 and R01 CA251339) and Memorial Sloan Kettering Cancer Center (P30 CA008748).

## CODE AVAILABILITY

Our algorithm is implemented in Python and available on GitHub at https://github.com/MSK-WANGX/BayesMAGPIE.

## Notes

### Competing Interest Statement

The authors have declared no competing interest.

## REFERENCES

Babur, O., Gönen, M., Aksoy, B. A., Schultz, N., Ciriello, G., Sander, C., and Demir, E. (2015). Systematic identification of cancer driving signaling pathways based on mutual exclusivity of genomic alterations. Genome biology 16, 1–10.

Bashashati, A., Haffari, G., Ding, J., et al. (2012). DriverNet: uncovering the impact of somatic driver mutations on transcriptional networks in cancer. Genome Biol 13, R124.

Cancer Genome Atlas, N. (2015). Genomic Classification of Cutaneous Melanoma. Cell 161, 1681–1696.

Canisius, S., Martens, J. W. M., and Wessels, L. F. A. (2016). A novel independence test for somatic alterations in cancer shows that biology drives mutual exclusivity but chance explains most co-occurrence. Genome biology 17, 1–17.

Ciriello, G., Cerami, E., Sander, C., and Schultz, N. (2012). Mutual exclusivity analysis identifies oncogenic network modules. Genome Res 22, 398–406.

Constantinescu, S., Szczurek, E., Mohammadi, P., Rahnenführer, J., and Beerenwinkel, N. (2016). TiMEx: a waiting time model for mutually exclusive cancer alterations. Bioinformatics 32, 968–975.

Ding, L., Getz, G., Wheeler, D. A., et al. (2008). Somatic mutations affect key pathways in lung adenocarcinoma. Nature 455, 1069–1075.

Fedrizzi, T., Ciani, Y., Lorenzin, F., Cantore, T., Gasperini, P., and Demichelis, F. (2021). Fast mutual exclusivity algorithm nominates potential synthetic lethal gene pairs through brute force matrix product computations. Computational and Structural Biotechnology Journal 19, 4394–4403.

Han, Y., Yang, J., Qian, X., et al. (2019). DriverML: a machine learning algorithm for identifying driver genes in cancer sequencing studies. Nucleic Acids Res 47, e45.

Hodis, E., Triglia, E. T., Kwon, J. Y. H., et al. (2022). Stepwise-edited, human melanoma models reveal mutations’ effect on tumor and microenvironment. Science 376, 474-+.

Hoffman, M. D., Blei, D. M., Wang, C., and Paisley, J. (2013). Stochastic variational inference. Journal of machine learning research.

Hou, J. P., and Ma, J. (2014). DawnRank: discovering personalized driver genes in cancer. Genome Med 6, 56.

Hu, Y., Yan, C., Chen, Q., and Meerzaman, D. (2021). gcMECM: graph clustering of mutual exclusivity of cancer mutations. BMC bioinformatics 22, 592.

Hua, X., Hyland, P. L., Huang, J., et al. (2016). MEGSA: A powerful and flexible framework for analyzing mutual exclusivity of tumor mutations. The American Journal of Human Genetics 98, 442–455.

Jia, P. L., Wang, Q., Chen, Q. X., Hutchinson, K. E., Pao, W., and Zhao, Z. M. (2014). MSEA: detection and quantification of mutation hotspots through mutation set enrichment analysis. Genome biology 15, 1–16.

Kim, Y. A., Madan, S., and Przytycka, T. M. (2017). WeSME: uncovering mutual exclusivity of cancer drivers and beyond. Bioinformatics 33, 814–821.

Kingma, D. P., and Ba, J. (2014). Adam: A method for stochastic optimization. *arXiv preprint arXiv:1412*.6980.

Lawrence, M. S., Stojanov, P., Polak, P., et al. (2013). Mutational heterogeneity in cancer and the search for new cancer-associated genes. Nature 499, 214–218.

Leiserson, M. D., Wu, H. T., Vandin, F., and Raphael, B. J. (2015). CoMEt: a statistical approach to identify combinations of mutually exclusive alterations in cancer. Genome Biol 16, 160.

Leiserson, M. D. M., Reyna, M. A., and Raphael, B. J. (2016). A weighted exact test for mutually exclusive mutations in cancer. Bioinformatics 32, 736–745.

Liu, S., Liu, J., Xie, Y., et al. (2021). MEScan: a powerful statistical framework for genome-scale mutual exclusivity analysis of cancer mutations. Bioinformatics 37, 1189–1197.

Mularoni, L., Sabarinathan, R., Deu-Pons, J., Gonzalez-Perez, A., and Lopez-Bigas, N. (2016). OncodriveFML: a general framework to identify coding and non-coding regions with cancer driver mutations. Genome Biol 17, 128.

Network, C. G. A. R. (2014). Comprehensive molecular profiling of lung adenocarcinoma. Nature 511, 543.

Paik, P. K., Felip, E., Veillon, R., et al. (2020). Tepotinib in non–small-cell lung cancer with MET exon 14 skipping mutations. New England Journal of Medicine 383, 931–943.

Pao, W., Wang, T. Y., Riely, G. J., et al. (2005). KRAS mutations and primary resistance of lung adenocarcinomas to gefitinib or erlotinib. PLoS Med 2, e17.

Reimand, J., and Bader, G. D. (2013). Systematic analysis of somatic mutations in phosphorylation signaling predicts novel cancer drivers. Mol Syst Biol 9, 637.

Sanchez-Vega, F., Mina, M., Armenia, J., et al. (2018). Oncogenic signaling pathways in the cancer genome atlas. Cell 173, 321–337. e310.

Shoushtari, A. N., Chatila, W. K., Arora, A., et al. (2021). Therapeutic Implications of Detecting MAPK-Activating Alterations in Cutaneous and Unknown Primary MelanomasMAP Kinase Drivers in Cutaneous and Unknown Primary Melanoma. Clinical Cancer Research 27, 2226–2235.

Szczurek, E., and Beerenwinkel, N. (2014). Modeling mutual exclusivity of cancer mutations. PLoS computational biology 10, e1003503.

Tamborero, D., Gonzalez-Perez, A., and Lopez-Bigas, N. (2013). OncodriveCLUST: exploiting the positional clustering of somatic mutations to identify cancer genes. Bioinformatics 29, 2238–2244.

Vandin, F., Upfal, E., and Raphael, B. J. (2012). De novo discovery of mutated driver pathways in cancer. Genome research 22, 375–385.

Wang, X., Kostrzewa, C., Reiner, A., Shen, R., and Begg, C. (2024). Adaptation of a mutual exclusivity framework to identify driver mutations within oncogenic pathways. The American Journal of Human Genetics 111, 227–241.

Wolf, J., Seto, T., Han, J.-Y., et al. (2020). Capmatinib in MET exon 14–mutated or MET-amplified non– small-cell lung cancer. New England Journal of Medicine 383, 944–957.

