## Supplemental Tables and Figures for "A Bayesian Approach for Identifying Driver Mutations within Oncogenic Pathways through Mutual Exclusivity"

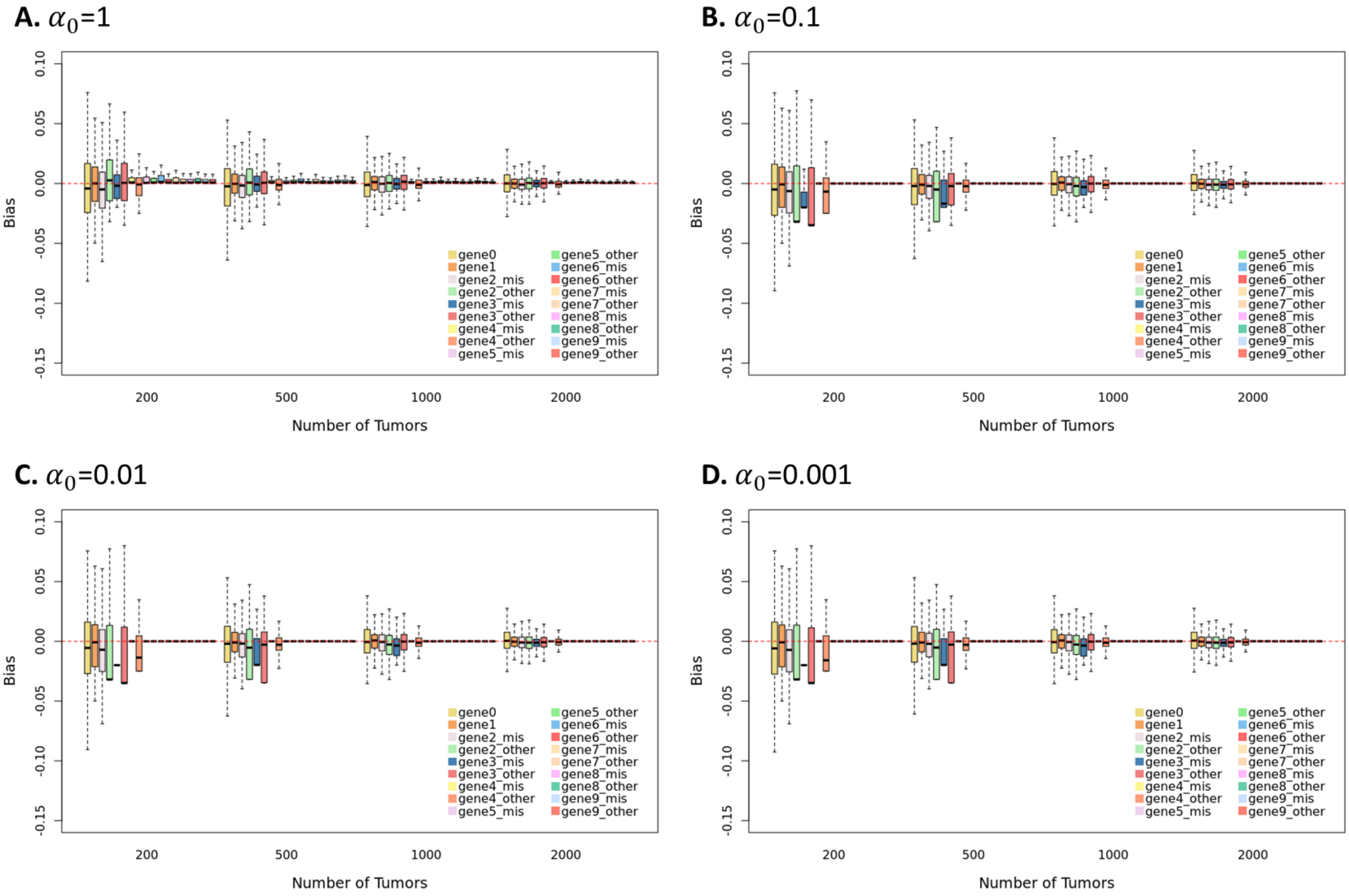


**Supplemental Figure 1** Boxplots Showing the Bias in the Estimated Driver Frequency of Each Gene from BayesMAGPIE under Varying Dirichlet Priors and Different Sample Sizes. A: $\alpha_0$=1; B: $\alpha_0$=0.1; C: $\alpha_0$=0.01; D: $\alpha_0$=0.001.

**Supplemental Table 1** Bias and standard deviation (SD) of the Estimated Driver Frequency of Each Gene from BayesMAGPIE under Varying Dirichlet Priors and Different Sample Sizes. A: $\alpha_0$=1; B: $\alpha_0$=0.1; C: $\alpha_0$=0.01; D: $\alpha_0$=0.001.

**1A.** $\alpha_0$=1

| | | | | N=200 | | N=500 | | N=1000 | | N=2000 | |
|---|---|---|---|---|---|---|---|---|---|---|---|
| | **Gene (Variant Type)** | **Simulated Mutation Freq** | **Simulated Driver Freq** | **Estimated Driver Freq** | **Bias (SD)** | **Estimated Driver Freq** | **Bias (SD)** | **Estimated Driver Freq** | **Bias (SD)** | **Estimated Driver Freq** | **Bias (SD)** |
| | **Gene 0** | 0.306 | 0.302 | 0.298 | -0.004 (0.032) | 0.299 | -0.003 (0.022) | 0.302 | 0 (0.015) | 0.302 | 0 (0.01) |
| | **Gene 1** | 0.084 | 0.050 | 0.05 | 0 (0.021) | 0.05 | 0 (0.012) | 0.051 | 0.001 (0.009) | 0.05 | 0 (0.006) |
| | **Gene 2** | | | | | | | | | | |
| | **Missense** | 0.076 | 0.069 | 0.064 | -0.005 (0.023) | 0.067 | -0.002 (0.014) | 0.068 | -0.001 (0.009) | 0.068 | -0.001 (0.007) |
| | **Other** | 0.083 | 0.032 | 0.036 | 0.004 (0.024) | 0.034 | 0.002 (0.016) | 0.033 | 0.001 (0.011) | 0.032 | 0 (0.007) |
| **Driver** | **Gene 3** | | | | | | | | | | |
| | **Missense** | 0.056 | 0.020 | 0.019 | -0.001 (0.014) | 0.02 | 0 (0.009) | 0.02 | 0 (0.007) | 0.02 | 0 (0.004) |
| | **Other** | 0.078 | 0.035 | 0.036 | 0.001 (0.021) | 0.036 | 0.001 (0.013) | 0.036 | 0.001 (0.01) | 0.035 | 0 (0.006) |
| | **Gene 4** | | | | | | | | | | |
| | **Missense** | 0.020 | 0 | 0.004 | 0.004 (0.006) | 0.002 | 0.002 (0.003) | 0.001 | 0.001 (0.002) | 0.001 | 0.001 (0.001) |
| | **Other** | 0.025 | 0.025 | 0.024 | -0.001 (0.012) | 0.024 | -0.001 (0.007) | 0.024 | -0.001 (0.005) | 0.025 | 0 (0.003) |
| | **Gene 5** | | | | | | | | | | |
| | **Missense** | 0.072 | 0 | 0.004 | 0.004 (0.007) | 0.002 | 0.002 (0.003) | 0.002 | 0.002 (0.002) | 0.001 | 0.001 (0.001) |
| | **Other** | 0.042 | 0 | 0.004 | 0.004 (0.006) | 0.003 | 0.003 (0.004) | 0.002 | 0.002 (0.002) | 0.001 | 0.001 (0.002) |
| | **Gene 6** | | | | | | | | | | |
| | **Missense** | 0.081 | 0 | 0.005 | 0.005 (0.008) | 0.003 | 0.003 (0.005) | 0.002 | 0.002 (0.003) | 0.002 | 0.002 (0.002) |
| | **Other** | 0.035 | 0 | 0.003 | 0.003 (0.005) | 0.002 | 0.002 (0.003) | 0.001 | 0.001 (0.002) | 0.001 | 0.001 (0.001) |
| | **Gene 7** | | | | | | | | | | |
| **Non-Driver** | **Missense** | 0.064 | 0 | 0.004 | 0.004 (0.007) | 0.003 | 0.003 (0.004) | 0.002 | 0.002 (0.002) | 0.001 | 0.001 (0.002) |
| | **Other** | 0.038 | 0 | 0.003 | 0.003 (0.006) | 0.002 | 0.002 (0.003) | 0.001 | 0.001 (0.002) | 0.001 | 0.001 (0.001) |
| | **Gene 8** | | | | | | | | | | |
| | **Missense** | 0.049 | 0 | 0.003 | 0.003 (0.006) | 0.002 | 0.002 (0.003) | 0.001 | 0.001 (0.002) | 0.001 | 0.001 (0.001) |
| | **Other** | 0.040 | 0 | 0.004 | 0.004 (0.006) | 0.002 | 0.002 (0.003) | 0.002 | 0.002 (0.002) | 0.001 | 0.001 (0.001) |
| | **Gene 9** | | | | | | | | | | |
| | **Missense** | 0.029 | 0 | 0.003 | 0.003 (0.005) | 0.002 | 0.002 (0.003) | 0.002 | 0.002 (0.002) | 0.001 | 0.001 (0.001) |
| | **Other** | 0.033 | 0 | 0.003 | 0.003 (0.006) | 0.002 | 0.002 (0.003) | 0.001 | 0.001 (0.002) | 0.001 | 0.001 (0.001) |

**1B.** $\alpha_0$=0.1

| | | | | N=200 | | N=500 | | N=1000 | | N=2000 | |
|---|---|---|---|---|---|---|---|---|---|---|---|
| | Gene (Variant Type) | Simulated Mutation Freq | Simulated Driver Freq | Estimated Driver Freq | Bias (SD) | Estimated Driver Freq | Bias (SD) | Estimated Driver Freq | Bias (SD) | Estimated Driver Freq | Bias (SD) |
| Driver | **Gene 0** | 0.306 | 0.302 | 0.296 | -0.006 (0.034) | 0.3 | -0.002 (0.021) | 0.303 | 0.001 (0.015) | 0.303 | 0.001 (0.01) |
| | **Gene 1** | 0.084 | 0.050 | 0.045 | -0.005 (0.027) | 0.049 | -0.001 (0.014) | 0.05 | 0 (0.009) | 0.05 | 0 (0.006) |
| | **Gene 2** | | | | | | | | | | |
| | **Missense** | 0.076 | 0.069 | 0.059 | -0.01 (0.029) | 0.066 | -0.003 (0.015) | 0.068 | -0.001 (0.009) | 0.068 | -0.001 (0.007) |
| | **Other** | 0.083 | 0.032 | 0.022 | -0.01 (0.03) | 0.026 | -0.006 (0.021) | 0.029 | -0.003 (0.014) | 0.031 | -0.001 (0.008) |
| | **Gene 3** | | | | | | | | | | |
| | **Missense** | 0.056 | 0.020 | 0.009 | -0.011 (0.016) | 0.012 | -0.008 (0.013) | 0.016 | -0.004 (0.009) | 0.019 | -0.001 (0.005) |
| | **Other** | 0.078 | 0.035 | 0.023 | -0.012 (0.028) | 0.029 | -0.006 (0.019) | 0.034 | -0.001 (0.012) | 0.034 | -0.001 (0.006) |
| | **Gene 4** | | | | | | | | | | |
| | **Missense** | 0.020 | 0 | 0 | 0 (0.001) | 0 | 0 (0.001) | 0 | 0 (0) | 0 | 0 (0) |
| | **Other** | 0.025 | 0.025 | 0.017 | -0.008 (0.017) | 0.022 | -0.003 (0.01) | 0.024 | -0.001 (0.006) | 0.025 | 0 (0.004) |
| Non-Driver | **Gene 5** | | | | | | | | | | |
| | **Missense** | 0.072 | 0 | 0 | 0 (0.003) | 0 | 0 (0.001) | 0 | 0 (0) | 0 | 0 (0) |
| | **Other** | 0.042 | 0 | 0 | 0 (0) | 0 | 0 (0) | 0 | 0 (0.001) | 0 | 0 (0) |
| | **Gene 6** | | | | | | | | | | |
| | **Missense** | 0.081 | 0 | 0 | 0 (0.001) | 0 | 0 (0.001) | 0 | 0 (0) | 0 | 0 (0) |
| | **Other** | 0.035 | 0 | 0 | 0 (0.001) | 0 | 0 (0.001) | 0 | 0 (0) | 0 | 0 (0) |
| | **Gene 7** | | | | | | | | | | |
| | **Missense** | 0.064 | 0 | 0 | 0 (0.003) | 0 | 0 (0.002) | 0 | 0 (0) | 0 | 0 (0) |
| | **Other** | 0.038 | 0 | 0 | 0 (0.002) | 0 | 0 (0.001) | 0 | 0 (0) | 0 | 0 (0) |
| | **Gene 8** | | | | | | | | | | |
| | **Missense** | 0.049 | 0 | 0 | 0 (0.001) | 0 | 0 (0) | 0 | 0 (0) | 0 | 0 (0) |
| | **Other** | 0.040 | 0 | 0 | 0 (0.001) | 0 | 0 (0) | 0 | 0 (0) | 0 | 0 (0) |
| | **Gene 9** | | | | | | | | | | |
| | **Missense** | 0.029 | 0 | 0 | 0 (0) | 0 | 0 (0) | 0 | 0 (0.001) | 0 | 0 (0) |
| | **Other** | 0.033 | 0 | 0 | 0 (0.003) | 0 | 0 (0) | 0 | 0 (0) | 0 | 0 (0) |

**1C.** $\alpha_0$=0.01

| | | | | N=200 | | N=500 | | N=1000 | | N=2000 | |
|---|---|---|---|---|---|---|---|---|---|---|---|
| | **Gene (Variant Type)** | **Simulated Mutation Freq** | **Simulated Driver Freq** | **Estimated Driver Freq** | **Bias (SD)** | **Estimated Driver Freq** | **Bias (SD)** | **Estimated Driver Freq** | **Bias (SD)** | **Estimated Driver Freq** | **Bias (SD)** |
| **Driver** | **Gene 0** | 0.306 | 0.302 | 0.296 | -0.006 (0.034) | 0.3 | -0.002 (0.021) | 0.303 | 0.001 (0.015) | 0.303 | 0.001 (0.01) |
| | **Gene 1** | 0.084 | 0.050 | 0.044 | -0.006 (0.027) | 0.049 | -0.001 (0.014) | 0.05 | 0 (0.009) | 0.05 | 0 (0.006) |
| | **Gene 2** | | | | | | | | | | |
| | **Missense** | 0.076 | 0.069 | 0.058 | -0.011 (0.03) | 0.066 | -0.003 (0.015) | 0.068 | -0.001 (0.009) | 0.068 | -0.001 (0.007) |
| | **Other** | 0.083 | 0.032 | 0.021 | -0.011 (0.03) | 0.025 | -0.007 (0.022) | 0.028 | -0.004 (0.015) | 0.031 | -0.001 (0.008) |
| | **Gene 3** | | | | | | | | | | |
| | **Missense** | 0.056 | 0.020 | 0.008 | -0.012 (0.016) | 0.011 | -0.009 (0.013) | 0.015 | -0.005 (0.01) | 0.019 | -0.001 (0.005) |
| | **Other** | 0.078 | 0.035 | 0.022 | -0.013 (0.028) | 0.028 | -0.007 (0.02) | 0.033 | -0.002 (0.012) | 0.034 | -0.001 (0.006) |
| | **Gene 4** | | | | | | | | | | |
| | **Missense** | 0.020 | 0 | 0 | 0 (0) | 0 | 0 (0.001) | 0 | 0 (0) | 0 | 0 (0) |
| | **Other** | 0.025 | 0.025 | 0.016 | -0.009 (0.017) | 0.022 | -0.003 (0.01) | 0.024 | -0.001 (0.006) | 0.025 | 0 (0.004) |
| **Non-Driver** | **Gene 5** | | | | | | | | | | |
| | **Missense** | 0.072 | 0 | 0 | 0 (0.002) | 0 | 0 (0.001) | 0 | 0 (0) | 0 | 0 (0) |
| | **Other** | 0.042 | 0 | 0 | 0 (0) | 0 | 0 (0) | 0 | 0 (0.001) | 0 | 0 (0) |
| | **Gene 6** | | | | | | | | | | |
| | **Missense** | 0.081 | 0 | 0 | 0 (0.001) | 0 | 0 (0) | 0 | 0 (0) | 0 | 0 (0) |
| | **Other** | 0.035 | 0 | 0 | 0 (0.001) | 0 | 0 (0.001) | 0 | 0 (0) | 0 | 0 (0) |
| | **Gene 7** | | | | | | | | | | |
| | **Missense** | 0.064 | 0 | 0 | 0 (0.002) | 0 | 0 (0.001) | 0 | 0 (0) | 0 | 0 (0) |
| | **Other** | 0.038 | 0 | 0 | 0 (0.001) | 0 | 0 (0) | 0 | 0 (0) | 0 | 0 (0) |
| | **Gene 8** | | | | | | | | | | |
| | **Missense** | 0.049 | 0 | 0 | 0 (0.001) | 0 | 0 (0) | 0 | 0 (0) | 0 | 0 (0) |
| | **Other** | 0.040 | 0 | 0 | 0 (0.001) | 0 | 0 (0) | 0 | 0 (0) | 0 | 0 (0) |
| | **Gene 9** | | | | | | | | | | |
| | **Missense** | 0.029 | 0 | 0 | 0 (0) | 0 | 0 (0) | 0 | 0 (0) | 0 | 0 (0) |
| | **Other** | 0.033 | 0 | 0 | 0 (0.003) | 0 | 0 (0) | 0 | 0 (0) | 0 | 0 (0) |

**1D.** $\alpha_0$=0.001

| | | | | N=200 | | N=500 | | N=1000 | | N=2000 | |
|---|---|---|---|---|---|---|---|---|---|---|---|
| | **Gene (Variant Type)** | **Simulated Mutation Freq** | **Simulated Driver Freq** | **Estimated Driver Freq** | **Bias (SD)** | **Estimated Driver Freq** | **Bias (SD)** | **Estimated Driver Freq** | **Bias (SD)** | **Estimated Driver Freq** | **Bias (SD)** |
| **Driver** | **Gene 0** | 0.306 | 0.302 | 0.296 | -0.006 (0.034) | 0.3 | -0.002 (0.021) | 0.303 | 0.001 (0.015) | 0.303 | 0.001 (0.01) |
| | **Gene 1** | 0.084 | 0.050 | 0.044 | -0.006 (0.027) | 0.049 | -0.001 (0.014) | 0.05 | 0 (0.009) | 0.05 | 0 (0.006) |
| | **Gene 2** | | | | | | | | | | |
| | **Missense** | 0.076 | 0.069 | 0.058 | -0.011 (0.03) | 0.066 | -0.003 (0.015) | 0.068 | -0.001 (0.009) | 0.068 | -0.001 (0.007) |
| | **Other** | 0.083 | 0.032 | 0.021 | -0.011 (0.03) | 0.025 | -0.007 (0.022) | 0.028 | -0.004 (0.015) | 0.031 | -0.001 (0.008) |
| | **Gene 3** | | | | | | | | | | |
| | **Missense** | 0.056 | 0.020 | 0.008 | -0.012 (0.015) | 0.011 | -0.009 (0.013) | 0.015 | -0.005 (0.01) | 0.018 | -0.002 (0.005) |
| | **Other** | 0.078 | 0.035 | 0.021 | -0.014 (0.028) | 0.028 | -0.007 (0.02) | 0.033 | -0.002 (0.012) | 0.034 | -0.001 (0.006) |
| | **Gene 4** | | | | | | | | | | |
| | **Missense** | 0.020 | 0 | 0 | 0 (0) | 0 | 0 (0.001) | 0 | 0 (0) | 0 | 0 (0) |
| | **Other** | 0.025 | 0.025 | 0.016 | -0.009 (0.017) | 0.021 | -0.004 (0.01) | 0.024 | -0.001 (0.006) | 0.025 | 0 (0.004) |
| **Non-Driver** | **Gene 5** | | | | | | | | | | |
| | **Missense** | 0.072 | 0 | 0 | 0 (0.002) | 0 | 0 (0.001) | 0 | 0 (0) | 0 | 0 (0) |
| | **Other** | 0.042 | 0 | 0 | 0 (0) | 0 | 0 (0) | 0 | 0 (0.001) | 0 | 0 (0) |
| | **Gene 6** | | | | | | | | | | |
| | **Missense** | 0.081 | 0 | 0 | 0 (0.001) | 0 | 0 (0) | 0 | 0 (0) | 0 | 0 (0) |
| | **Other** | 0.035 | 0 | 0 | 0 (0.001) | 0 | 0 (0.001) | 0 | 0 (0) | 0 | 0 (0) |
| | **Gene 7** | | | | | | | | | | |
| | **Missense** | 0.064 | 0 | 0 | 0 (0.003) | 0 | 0 (0.001) | 0 | 0 (0) | 0 | 0 (0) |
| | **Other** | 0.038 | 0 | 0 | 0 (0.001) | 0 | 0 (0) | 0 | 0 (0) | 0 | 0 (0) |
| | **Gene 8** | | | | | | | | | | |
| | **Missense** | 0.049 | 0 | 0 | 0 (0.001) | 0 | 0 (0) | 0 | 0 (0) | 0 | 0 (0) |
| | **Other** | 0.040 | 0 | 0 | 0 (0.001) | 0 | 0 (0) | 0 | 0 (0) | 0 | 0 (0) |
| | **Gene 9** | | | | | | | | | | |
| | **Missense** | 0.029 | 0 | 0 | 0 (0) | 0 | 0 (0) | 0 | 0 (0) | 0 | 0 (0) |
| | **Other** | 0.033 | 0 | 0 | 0 (0.003) | 0 | 0 (0) | 0 | 0 (0) | 0 | 0 (0) |
